# Tier-based standards for FAIR sequence data and metadata sharing in microbiome research

**DOI:** 10.1101/2025.02.06.636914

**Authors:** Lina Kim, Anton Lavrinienko, Zuzana Sebechlebska, Sven Stoltenberg, Nicholas A. Bokulich

**Affiliations:** Department of Health Sciences and Technology, ETH Zurich, Switzerland

**Author notes:** Corresponding author: Nicholas A. Bokulich. These authors contributed equally to this publication (Lina Kim, Anton Lavrinienko).

**Keywords:** microbiome, data availability statement, data sharing, meta-research, metadata

## Abstract

Microbiome research is a growing, data-driven field within the life sciences. While policies exist for sharing microbiome sequence data and using standardized metadata schemes, compliance among researchers varies. To promote open research data best practices in microbiome research and adjacent communities, we (1) propose two tiered badge systems to evaluate data/metadata sharing compliance, and (2) developed an automated evaluation tool to determine adherence to data reporting standards in publications with amplicon and metagenome sequence data. In a systematic evaluation of publications (n = 2929) spanning human gut microbiome research, and in three case studies of soil and gut microbiota used to manually validate the evaluation tool (n = 370), we found nearly half of publications do not meet minimum standards for sequence data availability. Moreover, poor standardization of metadata creates a high barrier to harmonization and cross-study comparison. Using this badge system and evaluation tool, our proof-of-concept work exposes the (i) ineffectiveness of sequence data availability statements, and (ii) lack of consistent metadata reports used for annotation of microbial data. We highlight the need for improved practices and infrastructure that reduce barriers to data submission and maximize reproducibility in microbiome research. We anticipate that our tiered badge framework will promote dialogue regarding data sharing practices and facilitate microbiome data reuse, supporting best practices that make microbiome data FAIR.

**Graphical Abstract:** 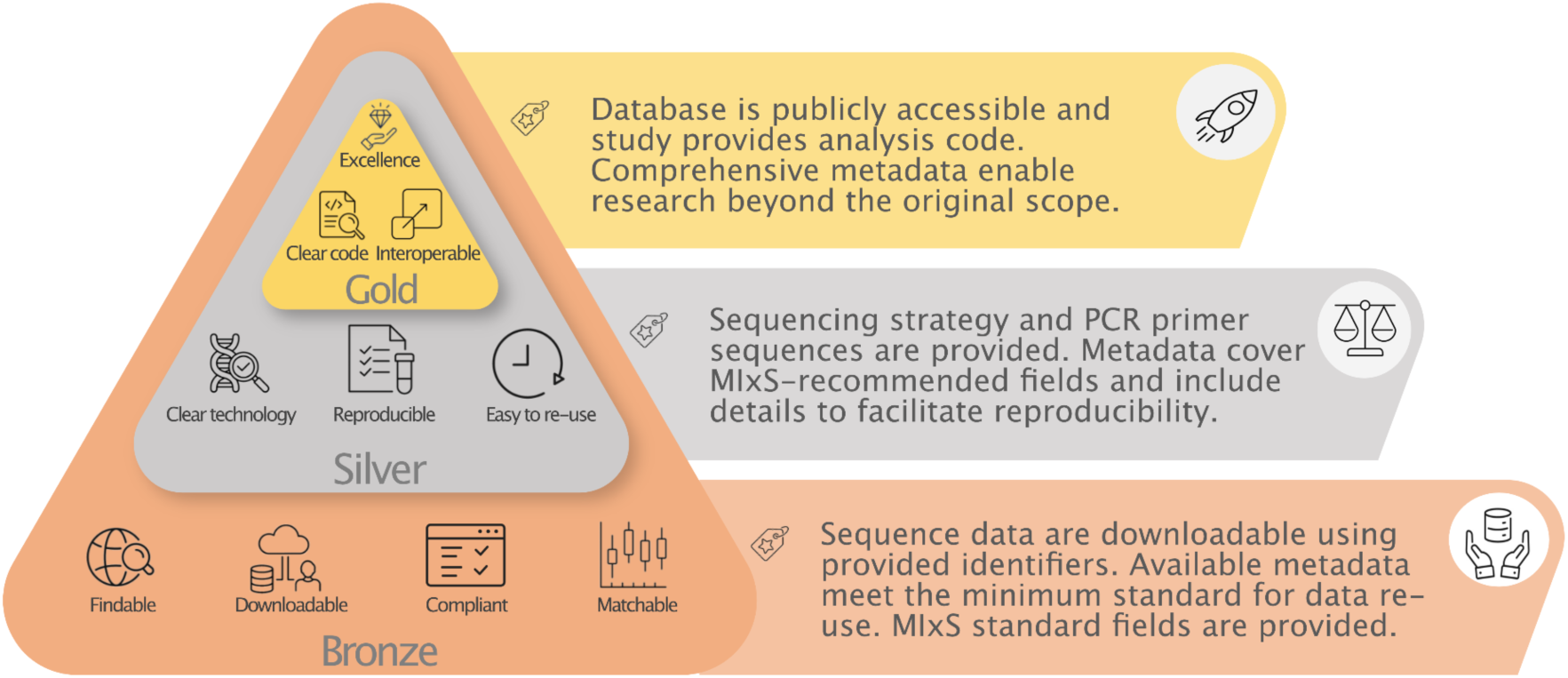

## Introduction

Reproducibility and data reuse are fundamental to advancing scientific discovery and innovation, yet both remain persistent challenges across the life sciences (1). Open research data (ORD) are publicly accessible, usable, and redistributable datasets. Ideally, ORD should also adhere to Findable, Accessible, Interoperable, and Reusable (FAIR) (2) principles and best practices that facilitate data reuse in research. Such openness is crucial in the life sciences, which are increasingly reliant on large-scale, high-dimensional datasets. Nearly half of all researchers use data generated by other scientists (3), and this number is likely to grow due to the increasing adoption of computational methods in all areas of research (4, 5) However, data reuse across research disciplines is hindered by subpar sharing practices including poor accessibility and interoperability, and particularly insufficient data and metadata reporting (10).

High-throughput DNA sequencing technologies have revolutionized research across all domains of the life sciences and over the past two decades have generated tremendous amounts of data (11). A prominent application of these technologies is the study of microbiomes: microbial communities, their components, products, interactions, and functional activities (12, 13) that play important roles in both host-associated and environmental ecosystems (6–9). Access to the raw sequence data and metadata is key to scientific transparency and innovative reuse, yet data accessibility (14) and metadata standardization issues (15) are salient in microbiome research.

Several data sharing standards already exist for depositing the raw microbiome sequence data, including theINSDC repositories (16, 17) as well as several microbiome-specific databases with integrated analytical workflows (e.g. Qiita (18), MGnify (19), and NMDC (20)). Similarly, various metadata standards were developed to establish a controlled vocabulary for sample collection, preparation, and data processing methods (21, 22). The Genomic Standards Consortium established the Minimum Information for any (x) Sequence standards (MIxS), with specific checklists for marker-gene (15) and metagenome data (23); these are currently supported by major sequence data repositories, including INSDC. Despite such institutional efforts, compliance with these data sharing standards among researchers is still limited (22).

Consequently, an increasing number of microbiome studies fail to meet standards and hamper the scientific community’s efforts toward open data sharing (24). Numerous studies either provide no accessible sequence data, or deposit the raw sequence data with insufficient or mismatched metadata that can not effectively be linked to the provided sequence data (25–27). This is a wider problem across numerous scientific disciplines, with publication data often remaining incomplete (28, 29). According to estimates, only a quarter of papers published in *Science* meet computational reproducibility (30) despite its policy requiring sharing of “all data necessary to understand, assess, and extend the conclusions of the manuscript”. Particularly frustrating is the refusal of authors to share published data despite reporting those as “available upon request” (31, 32). Indeed, the vast majority of researchers are not compliant with their own published data availability statements (DAS): of nearly 1,800 reviewed publications with DAS suggesting data availability upon “reasonable request”, a mere 7% of authors provided usable data (33).

Although depositing raw data prior to publication is now mandated by most journals and funding agencies, the key issue is that mechanisms for assessing and validating data reporting practices are largely missing (34). We propose a two-pronged solution to address these issues and promote reproducibility in the field of microbiome science: (i) tier-based standards to assess reproducibility of publications by evaluating data accessibility statements and metadata, and (ii) tools to programmatically determine adherence to these tiered standards, enabling evaluation pre-publication (e.g., during review) or post-publication (e.g., for selection during meta-analysis).

## Methods

### Tier-based standards for sequence data and metadata sharing

We propose two tiered badge systems to evaluate the openness of research data within the microbiome research community: one for sequence data reporting, and one for metadata reporting. These parallel standards are linked to provide effective data availability reporting for both nucleotide sequence data and associated metadata (Graphical Abstract, Tables 1-2). We consider in this framework the availability of next-generation sequencing (NGS)-based data, representing sequences comprising microbial communities (in the microbiome field, these are microbial amplicon and metagenome sequences). However, we anticipate that this framework can be extended and adopted by other fields within the life sciences relying on marker-gene (amplicon) data, such as for instance environmental DNA and diet metabarcoding research.

**Table 1.**
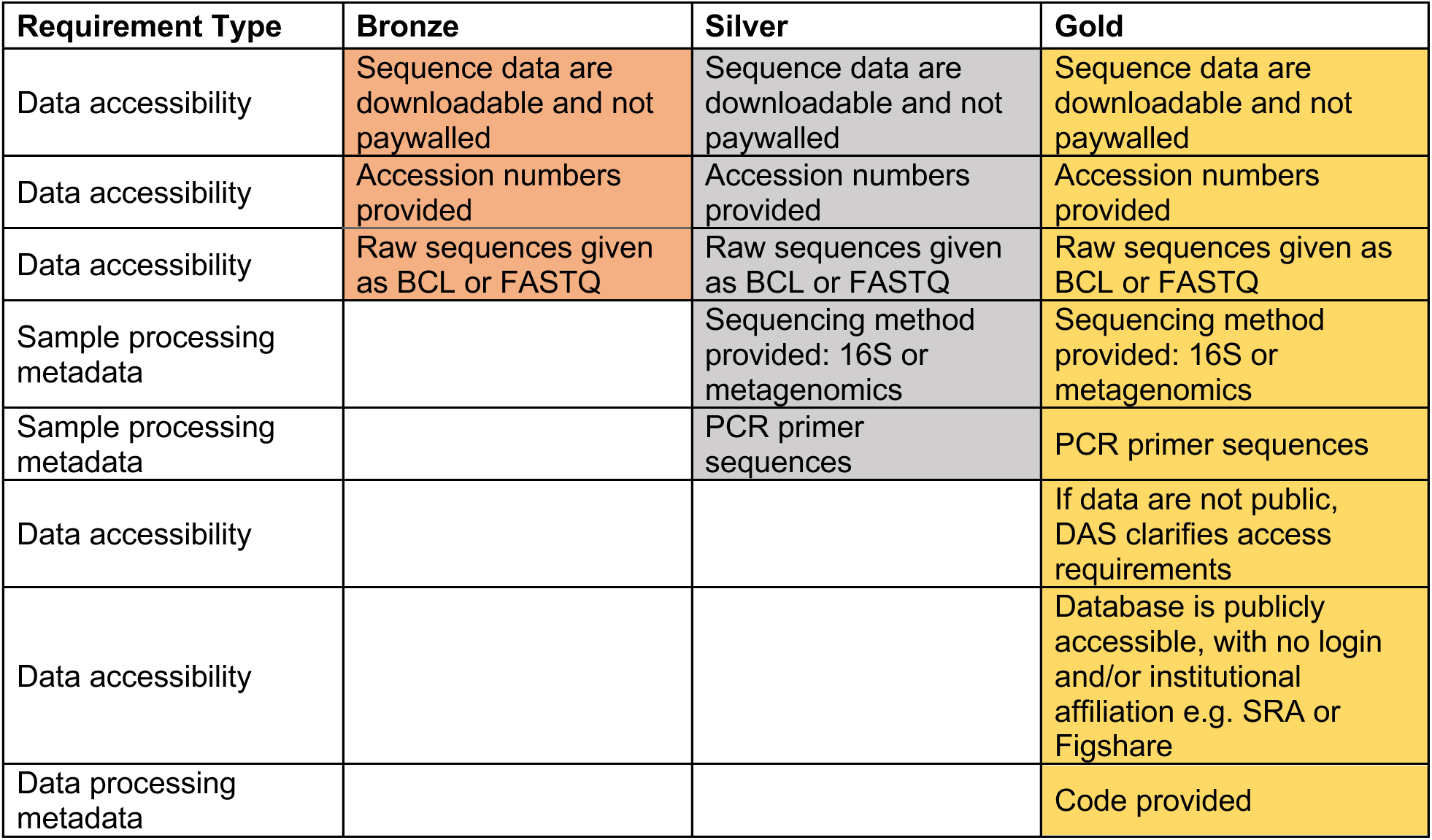
Requirements for nucleotide sequence data reporting. Bronze represents a minimal set of requirements for releasing open data, while the Gold tier represents both open and FAIR research data for easy reuse.

| Requirement Type | Bronze | Silver | Gold |
| --- | --- | --- | --- |
| Data accessibility | Sequence data are downloadable and not paywalled | Sequence data are downloadable and not paywalled | Sequence data are downloadable and not paywalled |
| Data accessibility | Accession numbers provided | Accession numbers provided | Accession numbers provided |
| Data accessibility | Raw sequences given as BCL or FASTQ | Raw sequences given as BCL or FASTQ | Raw sequences given as BCL or FASTQ |
| Sample processing metadata |  | Sequencing method provided: 16S or metagenomics | Sequencing method provided: 16S or metagenomics |
| Sample processing metadata |  | PCR primer sequences | PCR primer sequences |
| Data accessibility |  |  | If data are not public, DAS clarifies access requirements |
| Data accessibility |  |  | Database is publicly accessible, with no login and/or institutional affiliation e.g. SRA or Figshare |
| Data processing metadata |  |  | Code provided |

**Table 2.**
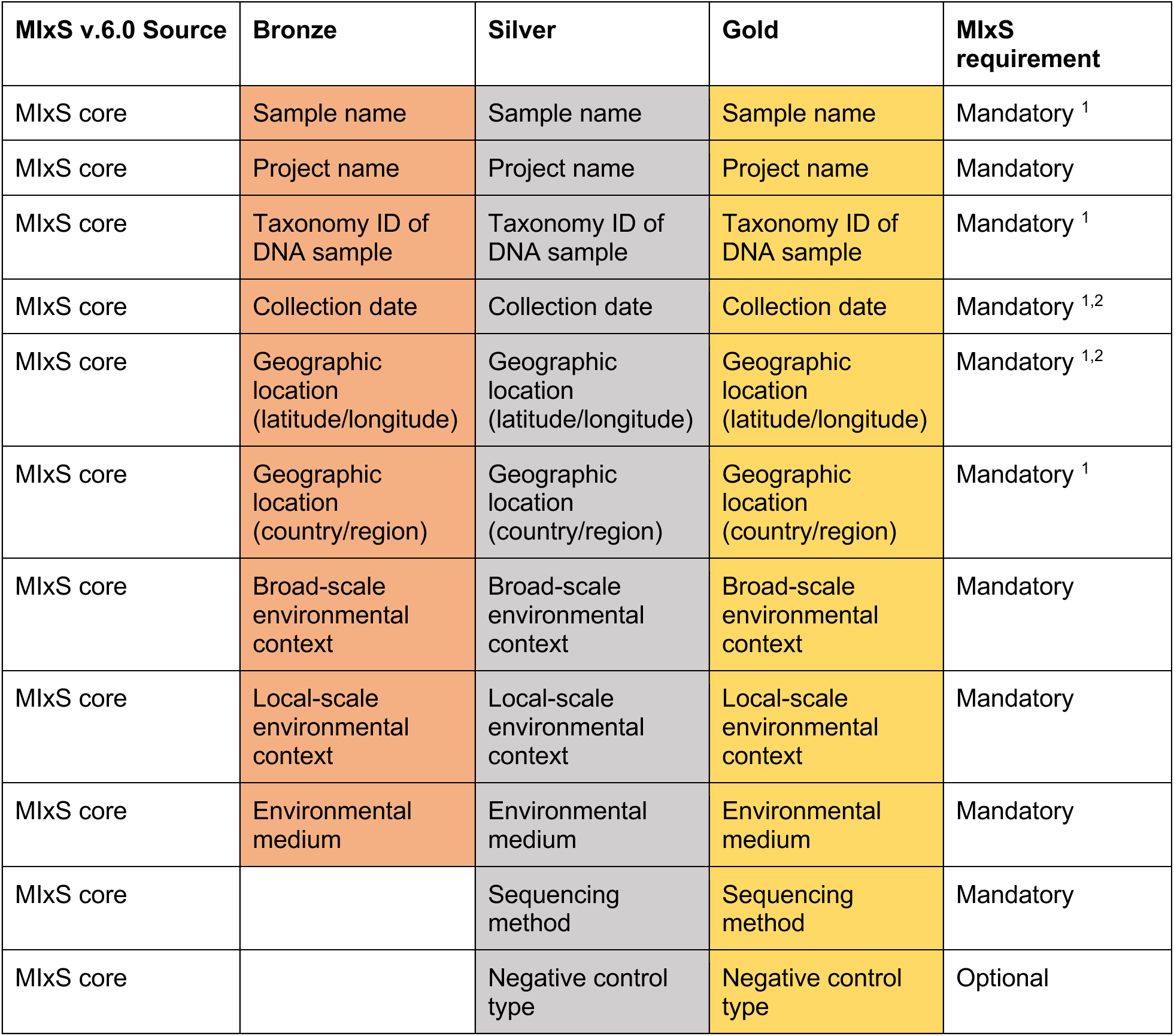
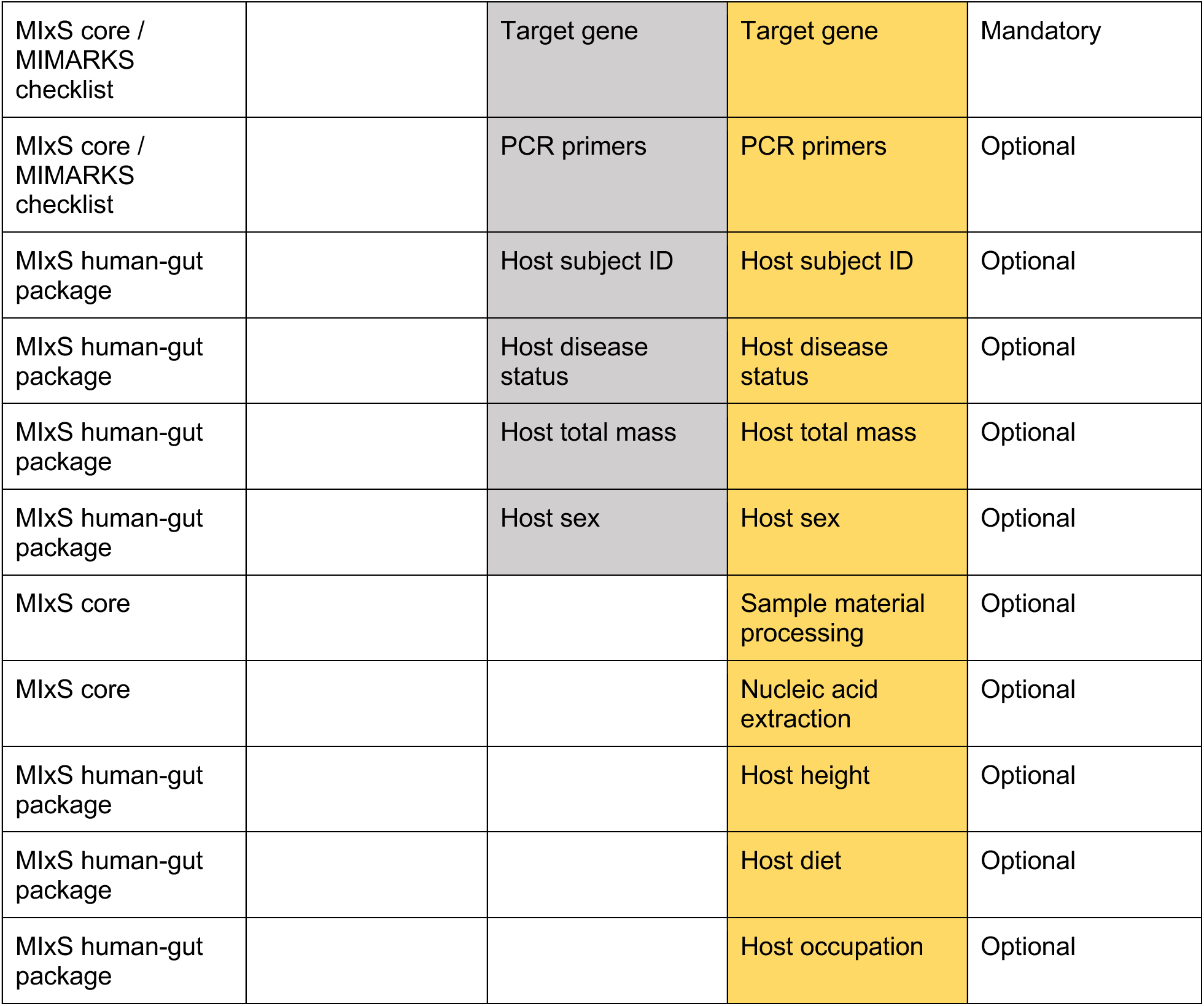
Example of requirements for metadata reporting. Bronze represents a minimal set of requirements for releasing open data, while the Gold tier represents both open and FAIR research data for easy reuse. Note that while in the example below the tiers do include the fields of a study that can be used to describe an experiment, or set of experiments, in an increasingly detailed manner (from Bronze to Gold), thus enabling data reuse, reproducibility, and eventually, research beyond the original scope, Silver and Gold are somewhat study-specific, i.e., these tiers do not assume the ever-increasing number of reported MIxS fields, but rather their quality and value for reproducibility and future data reuse. The MIxS-specific fields are based on the MIxS v6.0 release (https://www.ebi.ac.uk/ena/browser/view/ERC000015); ^1^ mandatory field in the NCBI metagenome or environmental package (https://submit.ncbi.nlm.nih.gov/biosample/template/), ^2^ mandatory in the ENA default sample checklist (https://www.ebi.ac.uk/ena/browser/view/ERC000011).

An ORD-compliant dataset requires that 1) the unprocessed sequence data and standardized metadata are available, ideally in standard formats (15), and 2) both are downloadable from a public repository or a database. The US National Center for Biotechnology Information (NCBI)’s Sequence Read Archive (SRA) and European Bioinformatics Institute (EBI)’s European Nucleotide Archive (ENA) are excellent examples of databases preserved in the public domain, providing “free, unrestricted and permanent access to the data” (35).

For a dataset to be FAIR (Findable, Accessible, Interoperable, and Reusable), additional properties are necessary beyond sequence data and metadata deposition. For example, to reproduce the publication’s analysis, knowledge of the conducted sequencing method (e.g., marker-gene vs. metagenomic sequencing) is necessary as bioinformatics analyses vary substantially and can be quite distinct (36) and/or require additional context. Notably, if the dataset was produced with marker-gene sequencing, primer sequences are necessary as primer choice affects sequence capture, database choice, and taxonomy identification (37, 38). Finally, full reproducibility and reuse of the raw dataset is only possible with the availability of the high-quality metadata and executable code used for the original analysis (39).

The badge system for sequence data and metadata availability standards ranks publications by integrating well accepted—but not yet universally adopted—standards:

**1. Bronze**: Available data meet the minimum standard for data reuse. To meet this standard, the publication includes basic details such as the accession ID for the associated sequence data, and the metadata that provides environmental context for given samples. With these, researchers are able to find and, when suitable for their purpose, use data from the original publication.
**2. Silver**: Facilitated reproducibility; this is a balance between comprehensiveness and feasibility for most researchers. This standard integrates more comprehensive metadata with additional fields such as recommended fields in the MixS checklists, and other domain-specific standards. Sequence data in this tier are not necessarily openly accessible, but the publication clarifies access requirements for reuse.
**3. Gold**: These data provide valuable information to address research questions beyond the original scope. With freely and publicly accessible sequence data, any researcher is able to use these data without barriers. And with additional optional metadata details from specific MIxS environmental packages, researchers are able to effectively characterize this dataset for deeper investigation.

### Automated badge evaluation tool

To automatically assess ORD reporting standards, we developed software to evaluate the openness of a given research publication. This software package is intended as a proof-of-concept for assessing the feasibility of automated screening of the nucleotide sequence data accessibility badge system described above (Table 1). The Python package MISHMASH (MIcrobiome Sequence and Metadata Availability Standards) automatically evaluates the openness of (1) nucleotide sequence data reporting, and (2) metadata reporting. It assesses given articles for the required reporting components listed in Tables 1-2, and uses this information to assign a badge (Bronze, Silver, or Gold). MISHMASH was developed as an open-source Python 3 package under the BSD-3-Clause license, and is available at https://github.com/bokulich-lab/mishmash.

With two main commands, users can evaluate data reporting in PubMed Central (PMC) publications. The command `assess_sequences` takes as input a space-separated list of PMC IDs or a file containing PMC IDs, where each ID corresponds to a single PMC publication. For each publication, the package returns a Sequence Accessibility Badge (Bronze, Silver, Gold, or “Cannot be determined”) based on the criteria in Table 1, and outputs the following components as a comma-separated values file (and indicates if these components were not detected, as transparent criteria for the badge assessment):

- INSDC database associated with the uploaded sequence data i.e. SRA, ENA, or DDBJ
- INSDC accession numbers corresponding to the sequence data uploaded to INSDC databases
- Number of sequence records (INSDC Runs) associated with the input article
- Primer sequences of amplified variable regions, if an amplicon-based study
- Code availability per study

The command `assess_metadata` retrieves metadata associated with a sequence record from an INSDC database. It returns the entirety of the metadata record as a comma-separated values file. If a particular GSC checklist or environmental package is used (e.g. MIMARKS or MIMS metagenome/environmental, human-gut), its name is returned as reported to the INSDC.

### Case Studies to test and validate evaluation tool

To assess the effectiveness of our standards and software, we manually validated the MISHMASH program and sequence data availability tiers on three case studies:

1. Case Study 1: a cross-section of human gut microbiome publications released between 2024 January 01 and 2024 June 30 (*n* = 85), and
2. Case Study 2: a list of human gut mycobiota publications from 2013 through 2023 (*n* = 199).
3. Case Study 3: a cross-section of soil microbiome publications released between 1 January 2023 and 31 December 2023 (n = 86).

Publications were screened on the NCBI PubMed database for peer-reviewed, original research papers with licenses allowing for full-text automated retrieval. The full PubMed search queries can be found with their corresponding PRISMA diagrams in the Supplemental Figures (Supplementary Figures 1-3). Each publication was manually evaluated for a badge assignment using the proposed tier criteria presented in Table 1.

For effective future evaluations of large datasets, we established a baseline accuracy rate for the automated badge prediction algorithm MISHMASH. By comparing manual badge assignments with programmatically-generated predictions for each of the 268 publications, we established a baseline accuracy rate for the automated badge prediction algorithm. Accuracy is reported as overall accuracy (sum of correct predictions over all predictions), macro-average F1, and overall weighted F1 score (40).

To investigate current metadata sharing practices, we manually examined 42 publications with INSDC-based datasets from Case Study 1. We extracted metadata associated with each article’s INSDC entry to evaluate badges for metadata reporting.

### Open field survey of human gut microbiome literature

To evaluate the openness and FAIRness of the microbiome research field, we compiled an open field survey dataset of human gut microbiome publications comprising 2,929 articles published between 2003-2023 (Figure 1). As in the above case studies, included publications were peer-reviewed, original research papers with licenses allowing for automated full-text retrieval. With the MISHMASH algorithm, we estimated the comprehensive state of data sharing from the past two decades of human gut microbiome literature.

**Figure 1.**
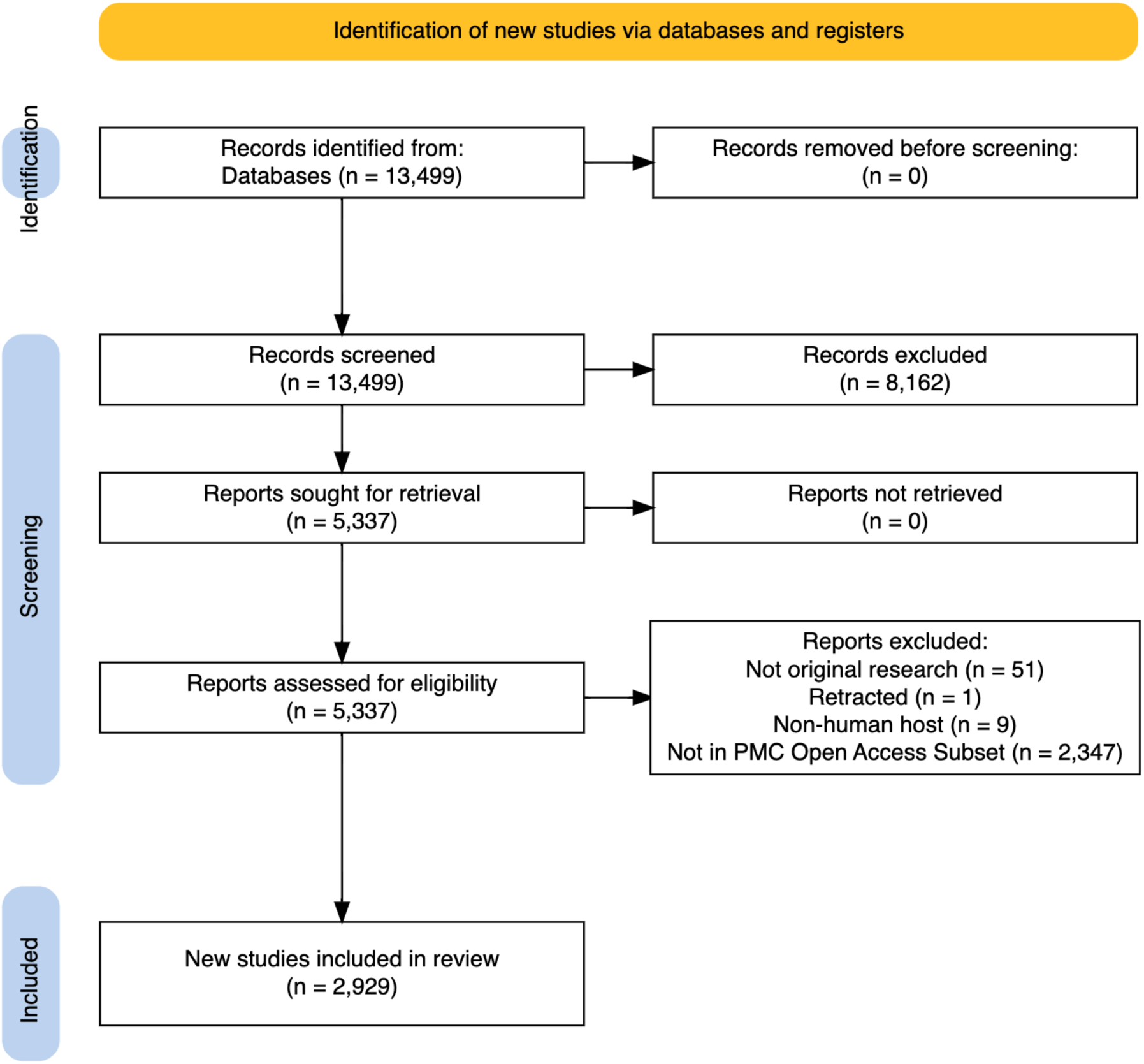
Overview of the workflow used for selection of publications for the open field survey. Records were removed during the initial screen (n = 8,162) for non-English text or non-human subjects based on MEDLINE indexing. The diagram was generated with PRISMA2020 (41).

## Results

### Manual error rate calculation demonstrates high accuracy

Using the studies in Case Study 1 and Case Study 2, we manually evaluated each publication’s badge assignment based on the tiered requirements established in Table 1; these manual assignments can be found in the Supplementary Tables. In both manually evaluated case studies, the most populated category was None, of publications that do not meet the minimum standards for open sequence data. The vast majority of remaining publications for both case studies were assigned the Silver badge, with similar proportions of both Bronze and Silver assignments.

As Case Study 2 spans ten years of publications, we additionally explored temporal patterns in data availability. The primary trend is of total frequency (Figure 3b): The number of annual publications steadily increases from one article in 2013 to 53 by the end of 2023. This is possibly explained by the decreasing cost of sequencing and growing interest in gut fungi in relation to human health. The proportion of publications without a badge assignment (Nones) also slowly decreases over time (Figure 3). We also investigated the potential influence of the article publisher on data openness. Of the 199 publications analyzed in Case Study 2, 153 (76.9%) were published in journals whose publishing groups included 10 or more publications from our list. Among these, data accessibility varies widely by publisher (Supplementary Figure 5): The proportion of published articles with no publicly available sequence data ranges from 18% to 44%.

Tested on the combined dataset of the aforementioned case studies, the automated badge predictor correctly classified a publication’s assigned badge at an overall accuracy of 84.0%. The discrepancy between the macro-average and weighted F1 scores (0.79 vs. 0.84) indicates varying error rates across different categories. The badge category Gold has the lowest support proportion (9.9%) with an F1 score of 0.62, while the Silver category with the highest support proportion (38.7%) has an F1 score of 0.82.

We also examined the accuracy rates for a more generalized classification of None vs. Badged (Bronze, Silver, and Gold) publications. This is relevant to cases in which researchers search for publications with minimally available sequence data. For this binary badge predictor, our automated algorithm evaluates with an overall accuracy of 97.2%.

### Metadata survey demonstrates poor harmonization

Of the 42 Case Study 1 publications with data uploaded to the INSDC, only 21 of these datasets used a MIxS metadata scheme. Those remaining submitted INSDC-required minimal metadata (e.g., SRA/ENA specific checklists); all datasets also included additional attributes relevant to each study.

While some metadata attributes such as Organism are complete for every INSDC-submitted sample, others are less commonly utilized. Sample names are submitted to the INSDC through seven different terms depending on the study and data submitter, such as “Sample Name” and “User_sample_ID”. Sample environmental contexts and microbiome host identifiers are likewise spread across variously named metadata attributes. In all, the union of metadata attributes from 42 studies totaled 220. Only 16 of these attributes were consistently filled across every study, and nearly 80% of metadata attributes had entries from fewer than eight publications (Figure 2). The median attribute included metadata from only a single study (2.4%), showcasing the immense sparsity of poorly labeled metadata.

**Figure 2.**
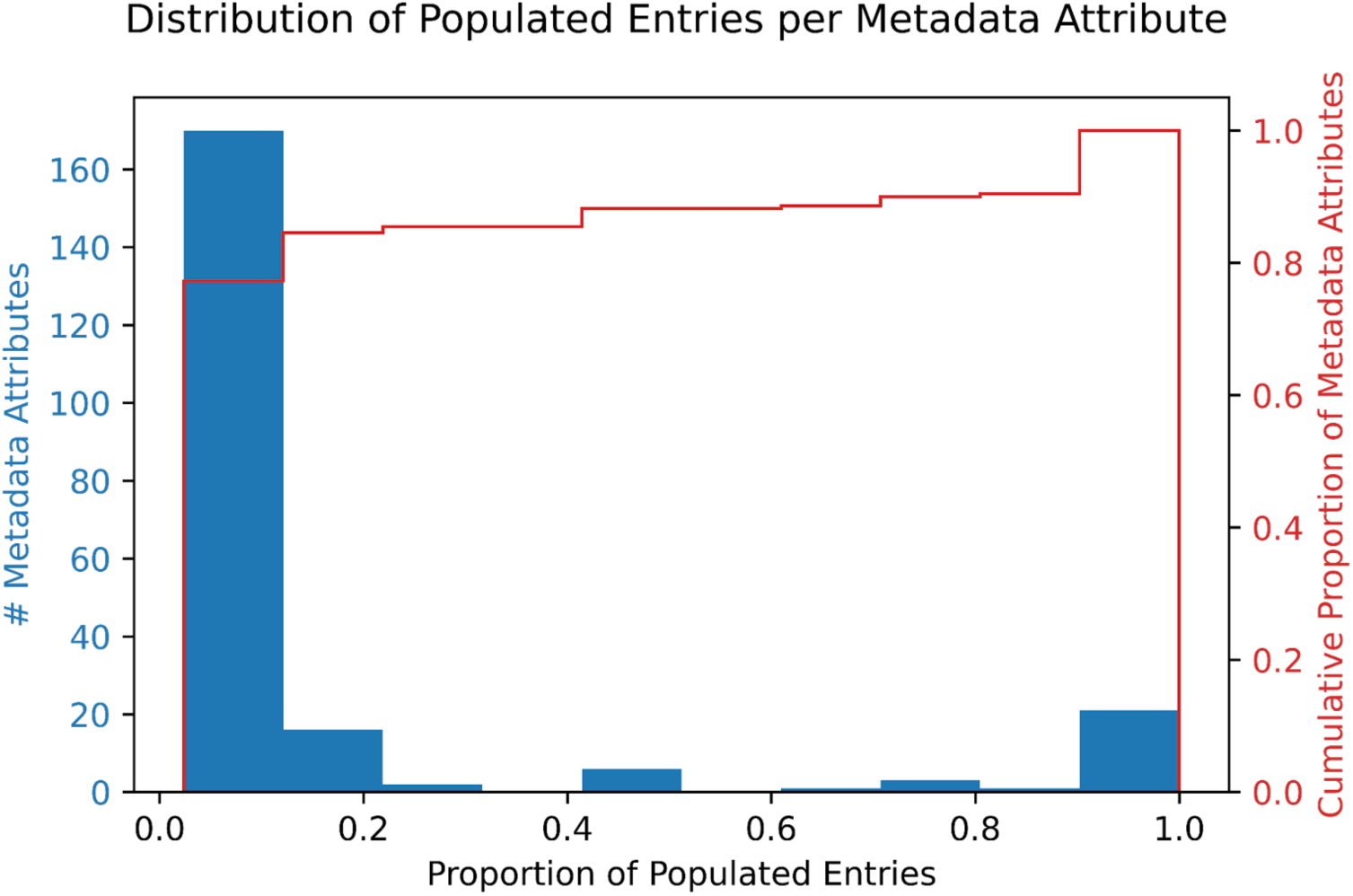
A histogram of metadata comprehensiveness across 42 studies and 220 metadata attributes. The proportion of populated entries indicates the percentage of publications with submitted metadata for a given attribute. For instance, 167/220 metadata attributes are populated at a rate of 10% or less, i.e. up to 90% of studies do not use or leave those metadata attributes blank.

### Open field survey and case studies show poor sequencing data accessibility

An open field survey of DAS found in microbiome research revealed three consistent trends: (i) nearly half of all publications (45.2%) did not meet the minimum standards for open sequence data (Nones), (ii) on the other extreme, only a few publications (8.0%) qualified for the Gold badge, and (iii) publication with the Silver badge (27.0%) outnumbered those with Bronze and Gold - consistent with our expectation that the Silver standard represents a balance between comprehensiveness and feasibility for many researchers. Our survey (Figure 3) yielded remarkably similar estimates to manual badge assignments for Case Studies 1 and 2, further reinforcing these findings and suggesting that such data reporting practices (i.e., high rates of missing sequence data) are consistent at the repository scale.

**Figure 3.**
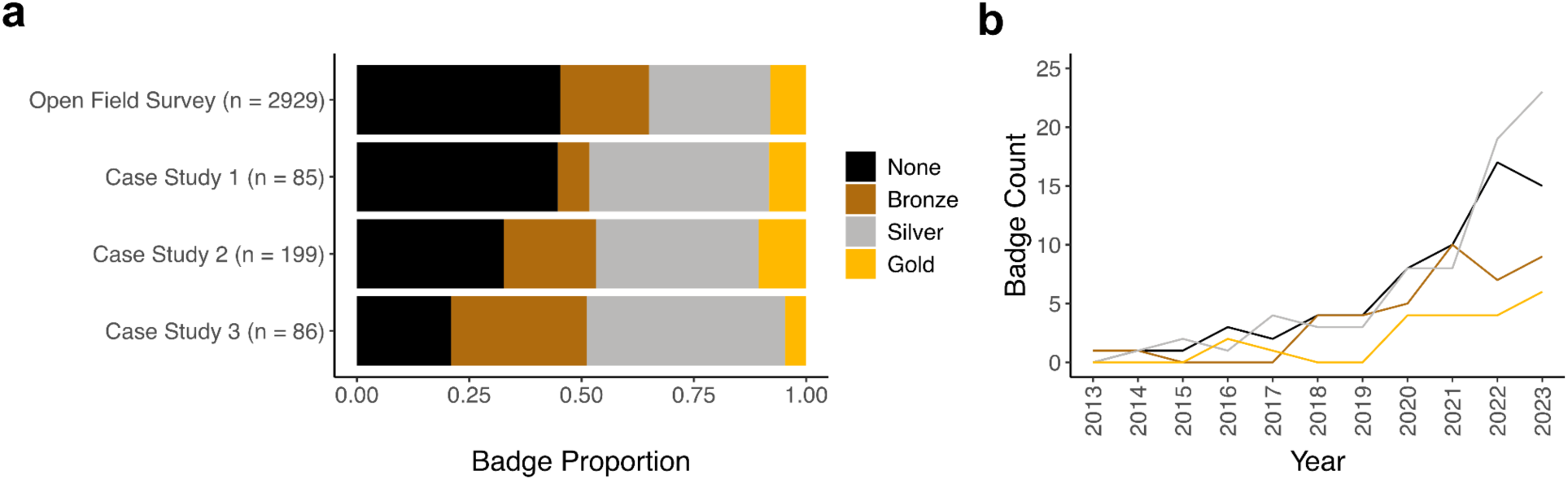
Trends of low open data compliance in microbiome literature. **a)** Predicted badge assignments for an open field survey of human gut-associated microbiome publications published from 2003 through 2023 (*n* = 2,929), and two smaller case studies with ground truth validation: badge assignments for human gut-associated microbiome publications published from January through June 2024 (*n =* 85), badge assignments for human fungal gut microbiome (mycobiome) publications published from 2013 through 2023 (*n =* 199), and badge assignments for soil microbiome publications published from January through December 2023 (*n =* 86). **b)** A temporal look at badge assignments for Case Study 2, by total frequency per year.

Interestingly, these trends likely reflect distinct user communities: researchers who simply share no data or report that data are “available upon request”, and researchers who exceed the minimum requirements, aiming for a higher standard of openness that balances availability with feasibility, as indicated by higher inclusion rates for Silver vs. Bronze badges in both case studies and our large-scale literature survey (Figure 3).

An evaluation of a completely separate research area — soil microbiota, as Case Study 3 — yielded similar proportions of data accessibility according to our badge system (Figure 3). In a pattern similar to those of human microbiome case studies, the proportion of Silver badges had the greatest plurality at 38/86 (44.2%) and Gold the smallest fraction at 4/86 (4.7%) of all publications. Case Studies 1 and 2 had Silver badge fractions of 40.0% and 35.5% respectively, and both had Gold-assigned papers as the smallest fractions at 8.2% and 10.4%. Case Study 3 had a far smaller proportion of None-badged publications than any of the previous case studies, at 20.9% compared with 44.7% (Case Study 1) and 33.3% (Case Study 2).

The error rates reported for our automated algorithm MISHMASH (Table 3) remain similar for this previously untested dataset. The overall accuracy of this four-category badge algorithm is 91.9%, greater than the 84.0% from Case Studies 1 and 2 combined (*n =* 268). The macro-average F1 and weighted F1 scores are both greater for Case Study 3, at 81.1% and 92.5% respectively. The overall accuracy for binary classification is comparable for Case Study 3, at 96.5% vs. 97.4% for Case Studies 1 and 2.

**Table 3.**
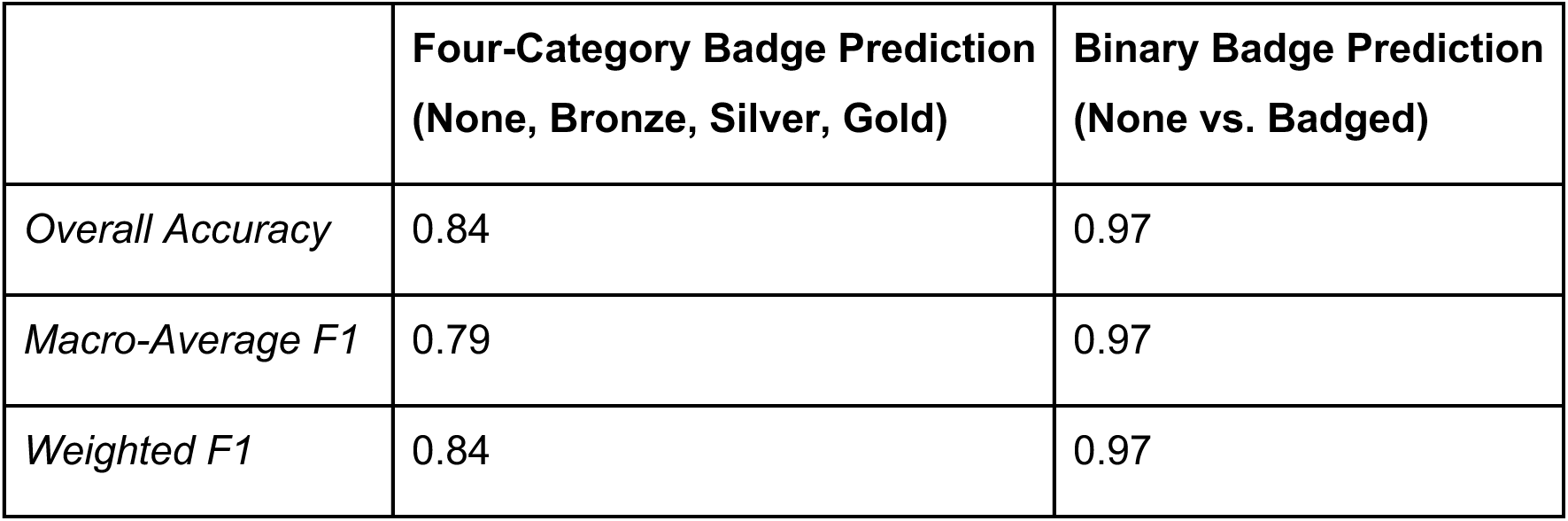
Accuracy rates for the automated badge prediction algorithm in establishing sequence data availability.

|  | **Four-Category Badge Prediction**<br>**(None, Bronze, Silver, Gold)** | **Binary Badge Prediction**<br>**(None vs. Badged)** |
| --- | --- | --- |
| *Overall Accuracy* | 0.84 | 0.97 |
| *Macro-Average F1* | 0.79 | 0.97 |
| *Weighted F1* | 0.84 | 0.97 |

### INSDC databases compose the bulk of reported sequence data

Microbiome publications upload their sequence data to numerous repositories. Among publications with associated sequence data (i.e. a badge of Bronze, Silver, or Gold), the majority (∼65%) utilized the NCBI Sequence Read Archive to store and share their data (Table 4). The European Nucleotide Archive, another INSDC database, is the second-most frequently used, representing ∼18% of total publications. The third INSDC database, the DNA Data Bank of Japan, falls at similar proportions to the China National Center for Bioinformation’s National Genomics Data Center (CNCB-NGDC) (42) used by ∼3% of publications screened.

**Table 4.**
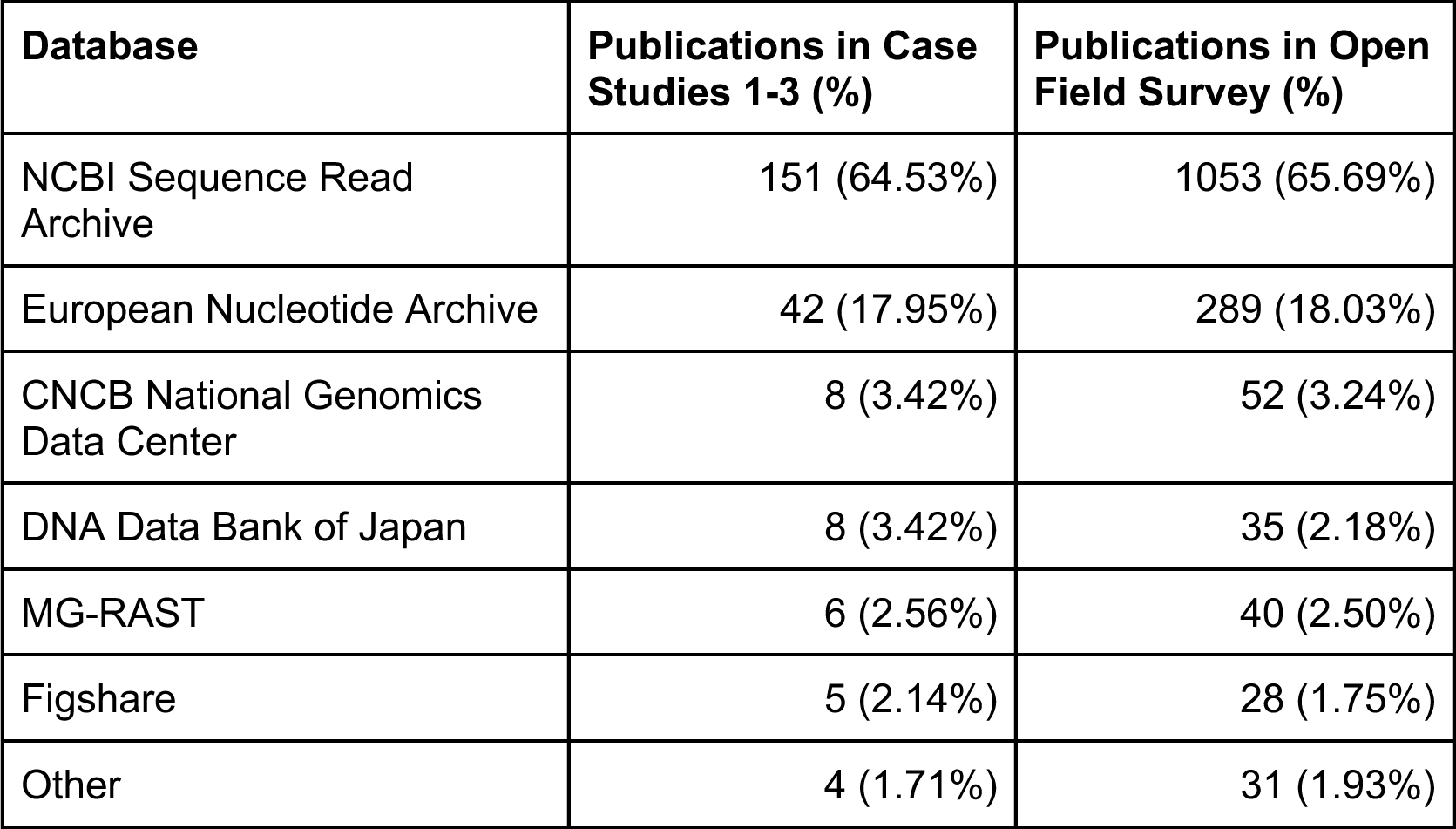
Distribution of utilized databases among microbiome publications. The “Other” category includes Zenodo, the European Genome-Phenome Archive, and the Chinese National GeneBank Sequence Archive. Publications uploading data to multiple databases comprise fewer than 5% of publications and were not included.

Much lower proportions of studies used the metagenome-specific platform MG-RAST, and Figshare (a surprising observation, as this repository is not specific to sequence data). Less frequently used are platforms such as Zenodo, the European Genome-Phenome Archive, and the Chinese National GeneBank Sequence Archive; these platforms each composed less than 1% of all publications each.

On the other hand, the distribution of badge assignments among utilized sequence repositories remained quite consistent. Publications from the Open Field Survey (*n =* 2929) aggregated similarly, with Bronze/Silver/Gold proportions among the most-frequently used archives (SRA and ENA) standing nearly identical (Figure 4).

**Figure 4.**
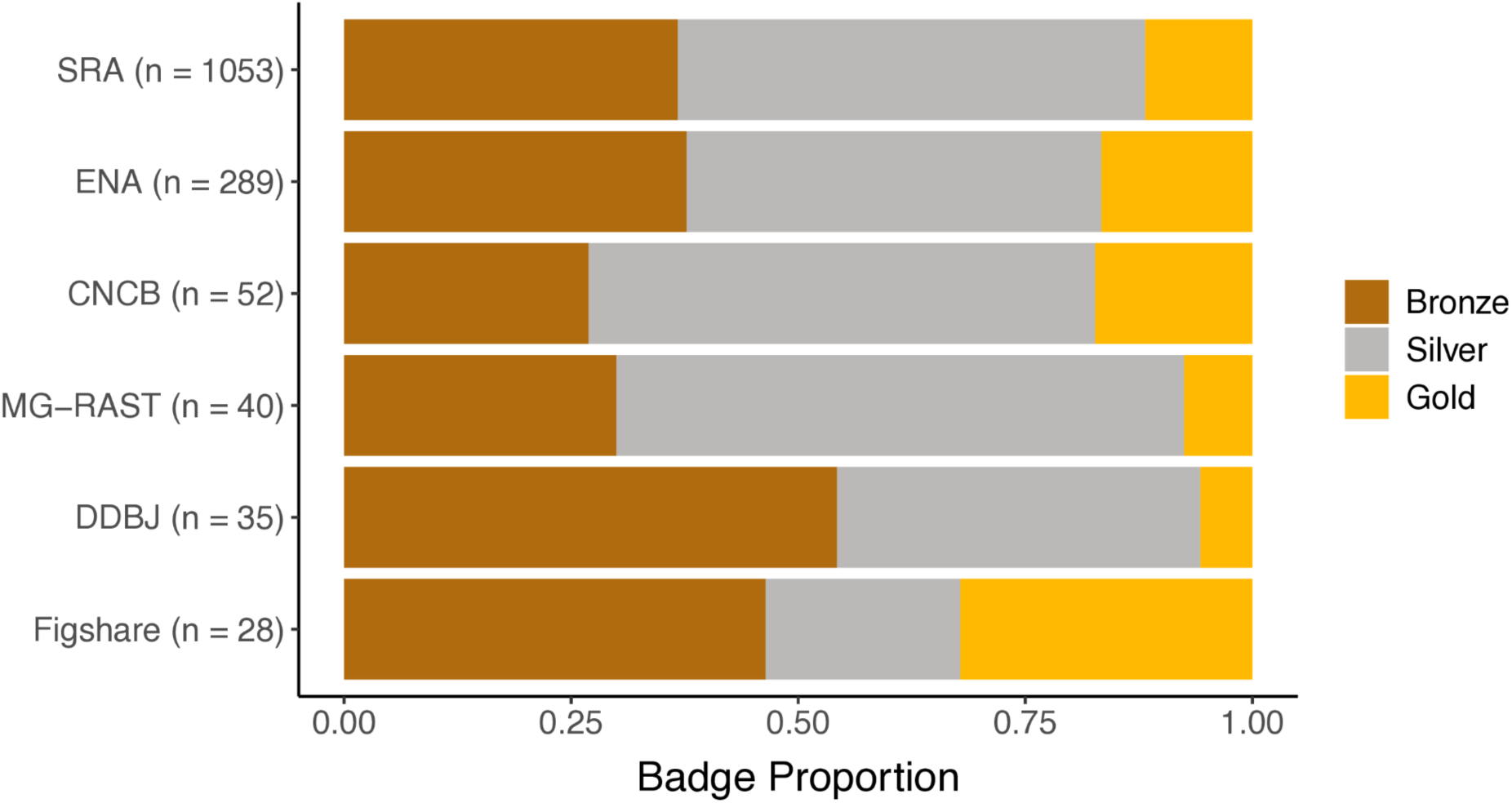
Distribution of badge assignments among utilized databases among papers from the open field survey.

## Discussion

### Systematic issues in current metadata sharing practices

Despite the critical role of metadata in microbiome research, standardized metadata are not commonly available, leaving microbiome datasets incomplete and limiting their utility (34). Indeed, most articles in our Case Study 1 did not qualify for a badge assignment when considering the reported metadata. This contrasts with the data availability statements for the same publications, where about half of the screened papers met eligibility for Bronze, Silver, or even Gold badges (Figure 3a). Metadata records are generally more complex and can be difficult to evaluate automatically using scraper tools. However, our implementation relies on a set of requirements for metadata reporting defined by the Genomics Standards Consortium’s MIxS checklists that should be standardized and machine-readable in a compliant report (15). Although adoption of the MIxS metadata standards is widespread across the microbiome community (22), metadata associated with most samples in INSDC databases remain poorly standardized, limiting the potential for microbiome data reuse. We found that the practice of adding non-standard metadata leads to considerable variation, making it challenging to harmonize metadata across studies.

More broadly, our findings highlight several systematic issues in current metadata collection and sharing practices. For example, several specific problems hinder efforts to evaluate existing metadata using the badge system alongside the DAS: (i) metadata often lack standardization and are reported without controlled vocabulary or standardized terminology, (ii) similar information is recorded in different fields across studies, (iii) information is reported, but the MIxS package requirements are applied incorrectly. In our Case Studies, the diversity of terms used to describe similar variables increased the dimensions of reported metadata, making it challenging to assess reporting practices automatically (Figure 2). More importantly, these same issues create barriers to data reuse, as it can be difficult or even impossible to harmonize such metadata records across studies without extensive manual inspection and curation. Similarly, a recent survey of public microbiome data identified over 3,000 different descriptors used to report common elements of sample metadata, such as those matching host age or sex (43). Such basic metadata associated with microbiome studies is also characterized by high levels of sparsity or missingness (e.g., >81.6% missing values for host age (43)). Although a number of resources and tools aimed at simplifying the annotation and formatting of sequence data and metadata exist (e.g., METAGENOTE (44)), their adoption is slow and these trends likely reflect the lack of centralized validation of standards or metadata quality during the submission process. As a result, even microbiome studies that share data in the INSDC databases (Bronze and above for DAS) often report only the minimally required metadata, which frequently falls short of minimum requirements for study-level reproducibility, lacks standardization, and contains free-text entries prone to inconsistencies and errors.

The misalignment between DAS and metadata reporting in terms of badge assignments in our Case Studies is concerning, as insufficient descriptive metadata directly limits the utility and future reuse of sequence data as they are inter-connected (45). Recognizing this issue, some initiatives are working on enriching microbiome data using text mining to extract relevant metadata terms from associated publications (46). Such metadata rescue efforts can help when basic structured metadata is missing (e.g., geographic annotation), but retrospective mapping of more complex metadata terms remains challenging. To this purpose, our proposed metadata badge system may serve as a qualitative guide for metadata reporting and harmonization in the microbiome field. As our current publication offers a prototype of badge tiers for human gut-associated metadata, we hope future iterations will adapt these requirements for different fields.

### Ensure quality by (self-)evaluating data sharing practices

We note that the high proportion (>45%) of None-badged publications with statements of “data available upon reasonable request” is concerning, as such data accessibility statements can include ostensible promises of data which often never materialize (33). Overall, these findings are consistent with proportions of data availability previously reported by Tedersoo et al. (2021) (14), in which an average of only 54.2% articles had accessible data prior to contact.

Temporal patterns in data availability reveal encouraging trends (Figure 3b). In the human fungal gut microbiome (mycobiome) studies, researchers have been adopting better data reporting practices at higher rates than in the mid-2010s (Supplementary Figure 4), likely due to stricter peer-review monitoring of data submission over the last decade and the growing number of initiatives that encourage data sharing by adding visibility to accessible data (1). Differences in data reporting guidelines (21) and the adoption rate of ORD standards (including variation in the peer-review process across journals) are also likely responsible for the apparent influence of a journal’s publisher on the distribution of assigned badges and data openness (Supplementary Figure 5).

As a single binary standard is insufficient to cover a wide range of cases, we propose a sliding scale of specific and measurable tiers that reflects usability and ease to other researchers (47). Our tiered “badge” system is designed to evaluate data reporting based on the level of information provided by authors (Graphical Abstract): Bronze-level contains the minimum required details for data accessibility and reuse, while Gold-ranked contributions allow for data access with little to no effort by users. Such a system provides coherent metrics to (self-)evaluate the quality of a dataset prior to investing time in reuse. In tandem with a user-friendly software for automated assessment, these unified standards and tier system will enable researchers, journals, and funding agencies to automatically assess compliance and quality of submitted datasets or publications, providing users with immediate, informative feedback (22, 48).

### Future directions

Our badge tier systems for sequencing data and metadata, and our associated prediction algorithm MISHMASH, are intended as proof-of-concepts. Their current iterations come with limitations, including a focus on the human gut microbiome in the given metadata badge standards. Future applications of metadata badge standards to non-human gut data should be adapted accordingly, possibly in the vein of MIxS standards applied to particular research fields. We note that due to heterogeneity in metadata reporting standards as described previously, the software MISHMASH does not currently provide metadata badge assessments. Future directions for improvement could include use of large language models to interpret and harmonize metadata.

While our prediction algorithm performs exceptionally well in establishing sequencing data availability (> 97%, Table 3), incomplete semantic processing necessitates limited manual checks and validation. Most assignment errors can be attributed to the difficulty in discriminating between badge tiers as opposed to between None vs. Badged publications, and can use continued improvement. In this initial proof-of-concept work, we were limited to databases with a robust API and thus were only able to evaluate open-access PMC publications available for web scraping (Figure 1); similarly, sequence data were only programmatically assessed if found in INSDC databases and certain common databases such as the Metagenomic Rapid Annotations using Subsystems Technology (MG-RAST) (49) or the Genome Sequence Archive of the Chinese National Genomics Data Center (GSA) (42). Although the PMC and INSDC do not encompass the entirety of scientific research, we expect our findings to be representative, as PMC and INSDC are among the largest and the most extensive research databases (Table 4). However, we hope that future iterations of our code will be able to automatically evaluate sequence data from less commonly used databases as well. We encourage the storage of sequence data in such databases rather than personal web pages due to issues with maintenance and “link rot” in the latter (50).

Our tier-based badge system promotes completely open data sharing, but an ethical issue arises in evaluating the accessibility of data locked under confidentiality agreements. Particularly with personally identifiable clinical data, patient confidentiality prevents completely public availability. Initiatives such as the European Genome-Phenome Archive (EGA) (51, 52) provide a workaround, as a secure archive released to authorized researchers. While data within the EGA is currently outside the scope of this proof-of-concept work, we would support a future integration to extend badging assessments to publications associated with the EGA and similar databases.

Additional training and coordinated educational campaigns will be needed to overcome perceived barriers to sharing data, which should start by building trust and increasing awareness regarding the value of open research data for individual scientists, wider scientific community, and society at large (53). Moreover, further user engagement will be needed to develop and extend this proof-of-concept resource to serve the specific needs of user communities across the microbiome research field (54). While here we focus on microbiome research, our tier-based standards and software approach are extensible and these principles can be easily adopted by life sciences fields relying on similar data types, such as environmental DNA analysis (55) and diet-based DNA metabarcoding (53). Additional training and coordinated educational campaigns will be needed to overcome perceived barriers to sharing data, which should start by building trust and increasing awareness regarding the value of open research data for individual scientists, wider scientific community, and society at large (50).

We hope that the release of our badge system and software will encourage self-reflection and greater engagement in FAIR practices in the microbiome community. Our vision for future implementation includes community outreach and engagement with stakeholders to encourage adoption of these or similar standards. We foresee the potential for integration with existing infrastructure to facilitate FAIR research awareness. Academic search engines such as the Swiss Library Service Platform, Web of Science, and Google Scholar include supplementary details with their search results; platforms such as these could include automated badge displays showcasing data availability with each hit to highlight commitment to FAIR principles. Additionally, these tiered data availability standards and software can be used by funding bodies or publishers to highlight and encourage data sharing practices among grantees or authors; for example, publishers who adopt these standards could integrate MISHMASH as an automated workflow to assign a badge during the publication process, and display badges alongside articles on the journal website. The ISME Journal already incorporates Open Data Badges (56, 57) to acknowledge and incentivize open science practices, demonstrating how a badge system could be implemented. Data sharing is required by a number of grants, institutions, and journals, but compliance can be challenging to manually check; this can be remedied with the use of automated software and quantifiable tiered standards.

## Conclusion

In parallel with other fields in the life sciences, microbiome research is an increasingly data-driven domain that relies on large-scale, high-dimensional datasets. Unrestricted access to sequence data and metadata is essential for scientific transparency and innovative reuse, and is therefore increasingly encouraged by the microbiome research community and required by many publishers and funding agencies. Despite the active development of data sharing standards and best practices in microbiome research, here we show that many studies still suffer from poor data accessibility and low-quality metadata, limiting efficient data reuse and constraining the potential benefits of meta-analyses and synthesis of published results. At least partly, this issue is due to the lack of mechanisms for assessing and validating data reporting practices and compliance with ORD reporting requirements, and lack of awareness of best practices among authors.

Preparing published sequence data and metadata for reuse is time-consuming and often inefficient due to the lack of validation methods. Here we propose tier-based standards and contribute an open-source software package intended for assessment of sequence data and metadata sharing practices in microbiome research by a diverse range of downstream users. We hope that these standards and software will help to promote better practices in microbiome research by serving multiple purposes: (i) enabling authors to self-assess compliance with FAIR principles, thereby increasing awareness and adoption of these standards in the field, and (ii) providing data repositories, publishers, funding agencies, and other stakeholders with a prototype model to automatically assess compliance of submitted datasets or publications. Moreover, this functionality will also facilitate meta-analysis as automated predictions can (iii) help users with rapid and accurate identification of studies with minimally available sequence data, and thus datasets potentially suitable for reuse.

Poor standardization of shared data remains a critical issue across the life sciences (1). We anticipate that evaluating data openness across publications will both promote data-sharing practices in the field and encourage greater community support and efforts toward FAIR ORD sharing and reporting. The sequence data and metadata badge systems we propose here are preliminary guides for reporting standards, to support efforts for improving FAIR literacy in the microbiome research domain and beyond.

## Data Availability

No new sequence data were generated for this study. PubMed Central and INSDC accession IDs used are listed in the Supplementary Tables. Code used for analysis and figure generation can be found at https://github.com/bokulich-publications/mishmash-fig.

## References

1. Price, E., Feyertag, F., Evans, T., Miskin, J., Mitrophanous, K. and Dikicioglu, D. (2024) What is the *real* value of omics data? Enhancing research outcomes and securing long-term data excellence. Nucleic Acids Res., 52, 12130–12140.

2. Wilkinson, M.D., Dumontier, M., Aalbersberg, Ij. J., Appleton, G., Axton, M., Baak, A., Blomberg, N., Boiten, J.-W., da Silva Santos, L.B., Bourne, P.E., et al. (2016) The FAIR Guiding Principles for scientific data management and stewardship. Sci. Data, 3, 160018.

3. Science Staff (2011) Challenges and Opportunities. Science, 331, 692–693.

4. Blei, D.M. and Smyth, P. (2017) Science and data science. Proc. Natl. Acad. Sci., 114, 8689– 8692.

5. Brus, L. (2020) The Rise of Computation. Nano Lett., 20, 801–802.

6. Thompson, L.R., Sanders, J.G., McDonald, D., Amir, A., Ladau, J., Locey, K.J., Prill, R.J., Tripathi, A., Gibbons, S.M., Ackermann, G., et al. (2017) A communal catalogue reveals Earth’s multiscale microbial diversity. Nature, 551, 457–463.

7. Cullin, N., Azevedo Antunes, C., Straussman, R., Stein-Thoeringer, C.K. and Elinav, E. (2021) Microbiome and cancer. Cancer Cell, 39, 1317–1341.

8. Averill, C., Anthony, M.A., Baldrian, P., Finkbeiner, F., Van Den Hoogen, J., Kiers, T., Kohout, P., Hirt, E., Smith, G.R. and Crowther, T.W. (2022) Defending Earth’s terrestrial microbiome. Nat. Microbiol., 7, 1717–1725.

9. Bokulich, N.A., Lewis, Z.T., Boundy-Mills, K. and Mills, D.A. (2016) A new perspective on microbial landscapes within food production. Curr. Opin. Biotechnol., 37, 182–189.

10. Wilson, S.L., Way, G.P., Bittremieux, W., Armache, J., Haendel, M.A. and Hoffman, M.M. (2021) Sharing biological data: why, when, and how. FEBS Lett., 595, 847–863.

11. Katz, K., Shutov, O., Lapoint, R., Kimelman, M., Brister, J.R. and O’Sullivan, C. (2022) The Sequence Read Archive: a decade more of explosive growth. Nucleic Acids Res., 50, D387–D390.

12. Blaser, M., Bork, P., Fraser, C., Knight, R. and Wang, J. (2013) The microbiome explored: recent insights and future challenges. Nat. Rev. Microbiol., 11, 213–217.

13. Eren, A.M. and Banfield, J.F. (2024) Modern microbiology: Embracing complexity through integration across scales. Cell, 187, 5151–5170.

14. Tedersoo, L., Küngas, R., Oras, E., Köster, K., Eenmaa, H., Leijen, Ä., Pedaste, M., Raju, M., Astapova, A., Lukner, H., et al. (2021) Data sharing practices and data availability upon request differ across scientific disciplines. Sci. Data, 8, 192.

15. Yilmaz, P., Kottmann, R., Field, D., Knight, R., Cole, J.R., Amaral-Zettler, L., Gilbert, J.A., Karsch-Mizrachi, I., Johnston, A., Cochrane, G., et al. (2011) Minimum information about a marker gene sequence (MIMARKS) and minimum information about any (x) sequence (MIxS) specifications. Nat. Biotechnol., 29, 415–420.

16. Arita, M., Karsch-Mizrachi, I. and Cochrane, G. (2021) The international nucleotide sequence database collaboration. Nucleic Acids Res., 49, D121–D124.

17. The international nucleotide sequence database collaboration (INSDC): enhancing global participation (2024) Nucleic Acids Res., 10.1093/nar/gkae1058.

18. Gonzalez, A., Navas-Molina, J.A., Kosciolek, T., McDonald, D., Vázquez-Baeza, Y., Ackermann, G., DeReus, J., Janssen, S., Swafford, A.D., Orchanian, S.B., et al. (2018) Qiita: rapid, web-enabled microbiome meta-analysis. Nat. Methods, 15, 796–798.

19. Richardson, L., Allen, B., Baldi, G., Beracochea, M., Bileschi, M.L., Burdett, T., Burgin, J., Caballero-Pérez, J., Cochrane, G., Colwell, L.J., et al. (2023) MGnify: the microbiome sequence data analysis resource in 2023. Nucleic Acids Res., 51, D753–D759.

20. The National Microbiome Data Collaborative Data Portal: an integrated multi-omics microbiome data resource (2022) Nucleic Acids Res., 10.1093/nar/gkab990.

21. Cernava, T., Rybakova, D., Buscot, F., Clavel, T., McHardy, A.C., Meyer, F., Meyer, F., Overmann, J., Stecher, B., Sessitsch, A., et al. (2022) Metadata harmonization–Standards are the key for a better usage of omics data for integrative microbiome analysis. *Environ*. Microbiome, 17, 33.

22. Vangay, P., Burgin, J., Johnston, A., Beck, K.L., Berrios, D.C., Blumberg, K., Canon, S., Chain, P., Chandonia, J.-M., Christianson, D., et al. (2021) Microbiome Metadata Standards: Report of the National Microbiome Data Collaborative’s Workshop and Follow-On Activities. mSystems, 6, e01194–20.

23. Consortium, T.G.S., Bowers, R.M., Kyrpides, N.C., Stepanauskas, R., Harmon-Smith, M., Doud, D., Reddy, T.B.K., Schulz, F., Jarett, J., Rivers, A.R., et al. (2017) Minimum information about a single amplified genome (MISAG) and a metagenome-assembled genome (MIMAG) of bacteria and archaea. Nat. Biotechnol., 35, 725–731.

24. Langille, M.G.I., Ravel, J. and Fricke, W.F. (2018) “Available upon request”: not good enough for microbiome data! Microbiome, 6, 8.

25. Broderick, D., Marsh, R., Waite, D., Pillarisetti, N., Chang, A.B. and Taylor, M.W. (2023) Realising respiratory microbiomic meta-analyses: time for a standardised framework. Microbiome, 11, 57.

26. Butcher, M.C., Short, B., Veena, C.L.R., Bradshaw, D., Pratten, J.R., McLean, W., Shaban, S.M.A., Ramage, G. and Delaney, C. (2022) Meta-analysis of caries microbiome studies can improve upon disease prediction outcomes. APMIS, 130, 763–777.

27. Data availability is the golden rule in research (2024) >Nat. Microbiol., 9, 879–879.

28. Tenopir, C., Dalton, E.D., Allard, S., Frame, M., Pjesivac, I., Birch, B., Pollock, D. and Dorsett, K. (2015) Changes in Data Sharing and Data Reuse Practices and Perceptions among Scientists Worldwide. PLOS ONE, 10, e0134826.

29. Roche, D.G., Kruuk, L.E.B., Lanfear, R. and Binning, S.A. (2015) Public Data Archiving in Ecology and Evolution: How Well Are We Doing? PLOS Biol., 13, e1002295.

30. Stodden, V., Seiler, J. and Ma, Z. (2018) An empirical analysis of journal policy effectiveness for computational reproducibility. Proc. Natl. Acad. Sci., 115, 2584–2589.

31. Borghi, J.A. and Gulick, A.E. (2018) Data management and sharing in neuroimaging: Practices and perceptions of MRI researchers. PLOS ONE, 13, 0200562.

32. Ventresca, M., Schünemann, H.J., Macbeth, F., Clarke, M., Thabane, L., Griffiths, G., Noble, S., Garcia, D., Marcucci, M., Iorio, A., et al. (2020) Obtaining and managing data sets for individual participant data meta-analysis: scoping review and practical guide. BMC Med. Res. Methodol., 20, 113.

33. Gabelica, M., Bojčić, R. and Puljak, L. (2022) Many researchers were not compliant with their published data sharing statement: a mixed-methods study. J. Clin. Epidemiol., 150, 33– 41.

34. Huttenhower, C., Finn, R.D. and McHardy, A.C. (2023) Challenges and opportunities in sharing microbiome data and analyses. Nat. Microbiol., 10.1038/s41564-023-01484-x.

35. Leinonen, R., Sugawara, H., Shumway, M., and on behalf of the International Nucleotide Sequence Database Collaboration (2011) The Sequence Read Archive. Nucleic Acids Res., 39, D19–D21.

36. Liu, Y.-X., Qin, Y., Chen, T., Lu, M., Qian, X., Guo, X. and Bai, Y. (2021) A practical guide to amplicon and metagenomic analysis of microbiome data. Protein Cell, 12, 315–330.

37. Fadeev, E., Cardozo-Mino, M.G., Rapp, J.Z., Bienhold, C., Salter, I., Salman-Carvalho, V., Molari, M., Tegetmeyer, H.E., Buttigieg, P.L. and Boetius, A. (2021) Comparison of Two 16S rRNA Primers (V3–V4 and V4–V5) for Studies of Arctic Microbial Communities. Front. Microbiol., 12, 637526.

38. Na, H.S., Song, Y., Yu, Y. and Chung, J. (2023) Comparative Analysis of Primers Used for 16S rRNA Gene Sequencing in Oral Microbiome Studies. Methods Protoc., 6, 71.

39. Peng, R.D. (2011) Reproducible Research in Computational Science. Science, 334, 1226–1227.

40. Powers, D.M.W. (2008) Evaluation: From Precision, Recall, and F-Factor to ROC, Informedness, Markedness & Correlation. In Technical Report SIE-07-001.

41. Haddaway, N.R., Page, M.J., Pritchard, C.C. and McGuinness, L.A. (2022) *PRISMA2020* : An R package and Shiny app for producing PRISMA 2020-compliant flow diagrams, with interactivity for optimised digital transparency and Open Synthesis. Campbell Syst. Rev., 18, e1230.

42. CNCB-NGDC Members and Partners, Bai, X., Bao, Y., Bei, S., Bu, C., Cao, R., Cao, Y., Cen, H., Chao, J., Chen, F., et al. (2024) Database Resources of the National Genomics Data Center, China National Center for Bioinformation in 2024. Nucleic Acids Res., 52, D18– D32.

43. Rosenfeld, G., Angelova, A., Shin, C., Quinones, M. and Hurt, D. (2021) Current challenges in microbiome metadata collection. 10.1101/2021.05.05.442781.

44. Quiñones, M., Liou, D.T., Shyu, C., Kim, W., Vujkovic-Cvijin, I., Belkaid, Y. and Hurt, D.E. (2020) “METAGENOTE: a simplified web platform for metadata annotation of genomic samples and streamlined submission to NCBI’s sequence read archive”. BMC Bioinformatics, 21, 378.

45. Huttenhower, C., Finn, R.D. and McHardy, A.C. (2023) Challenges and opportunities in sharing microbiome data and analyses. Nat. Microbiol., 8, 1960–1970.

46. Mitchell, A.L., Almeida, A., Beracochea, M., Boland, M., Burgin, J., Cochrane, G., Crusoe, M.R., Kale, V., Potter, S.C., Richardson, L.J., et al. (2019) MGnify: the microbiome analysis resource in 2020. Nucleic Acids Res., 10.1093/nar/gkz1035.

47. Blei, D.M. and Smyth, P. (2017) Science and data science. Proc. Natl. Acad. Sci., 114, 8689– 8692.

48. Schloss, P.D. (2018) Identifying and Overcoming Threats to Reproducibility, Replicability, Robustness, and Generalizability in Microbiome Research. mBio, 9, e00525–18.

49. Keegan, K.P., Glass, E.M. and Meyer, F. (2016) MG-RAST, a Metagenomics Service for Analysis of Microbial Community Structure and Function. In Martin, F., Uroz, S. (eds), Microbial Environmental Genomics (MEG), Methods in Molecular Biology. Springer New York, New York, NY, Vol. 1399, pp. 207–233.

50. Schloss, P.D. (2018) Identifying and Overcoming Threats to Reproducibility, Replicability, Robustness, and Generalizability in Microbiome Research. mBio, 9.

51. Freeberg, M.A., Fromont, L.A., D’Altri, T., Romero, A.F., Ciges, J.I., Jene, A., Kerry, G., Moldes, M., Ariosa, R., Bahena, S., et al. (2022) The European Genome-phenome Archive in 2021. Nucleic Acids Res., 50, D980–D987.

52. Lappalainen, I., Almeida-King, J., Kumanduri, V., Senf, A., Spalding, J.D., ur-Rehman, S., Saunders, G., Kandasamy, J., Caccamo, M., Leinonen, R., et al. (2015) The European Genome-phenome Archive of human data consented for biomedical research. Nat. Genet., 47, 692–695.

53. De Sousa, L.L., Silva, S.M. and Xavier, R. (2019) DNA metabarcoding in diet studies: Unveiling ecological aspects in aquatic and terrestrial ecosystems. *Environ*. DNA, 1, 199–214.

54. Gomes, D.G.E., Pottier, P., Crystal-Ornelas, R., Hudgins, E.J., Foroughirad, V., Sánchez-Reyes, L.L., Turba, R., Martinez, P.A., Moreau, D., Bertram, M.G., et al. (2022) Why don’t we share data and code? Perceived barriers and benefits to public archiving practices. Proc. R. Soc. B Biol. Sci., 289, 20221113.

55. Shokralla, S., Spall, J.L., Gibson, J.F. and Hajibabaei, M. (2012) Next-generation sequencing technologies for environmental DNA research. Mol. Ecol., 21, 1794–1805.

56. The ISME Journal Instructions to Authors. Oxf. Acad. Press ISME J.

57. Kidwell, M.C., Lazarević, L.B., Baranski, E., Hardwicke, T.E., Piechowski, S., Falkenberg, L.-S., Kennett, C., Slowik, A., Sonnleitner, C., Hess-Holden, C., et al. (2016) Badges to Acknowledge Open Practices: A Simple, Low-Cost, Effective Method for Increasing Transparency. PLOS Biol., 14, e1002456.

