## Supplemental Figures for "Tier-based standards for FAIR sequence data and metadata sharing in microbiome research"

### Supplementary Figures

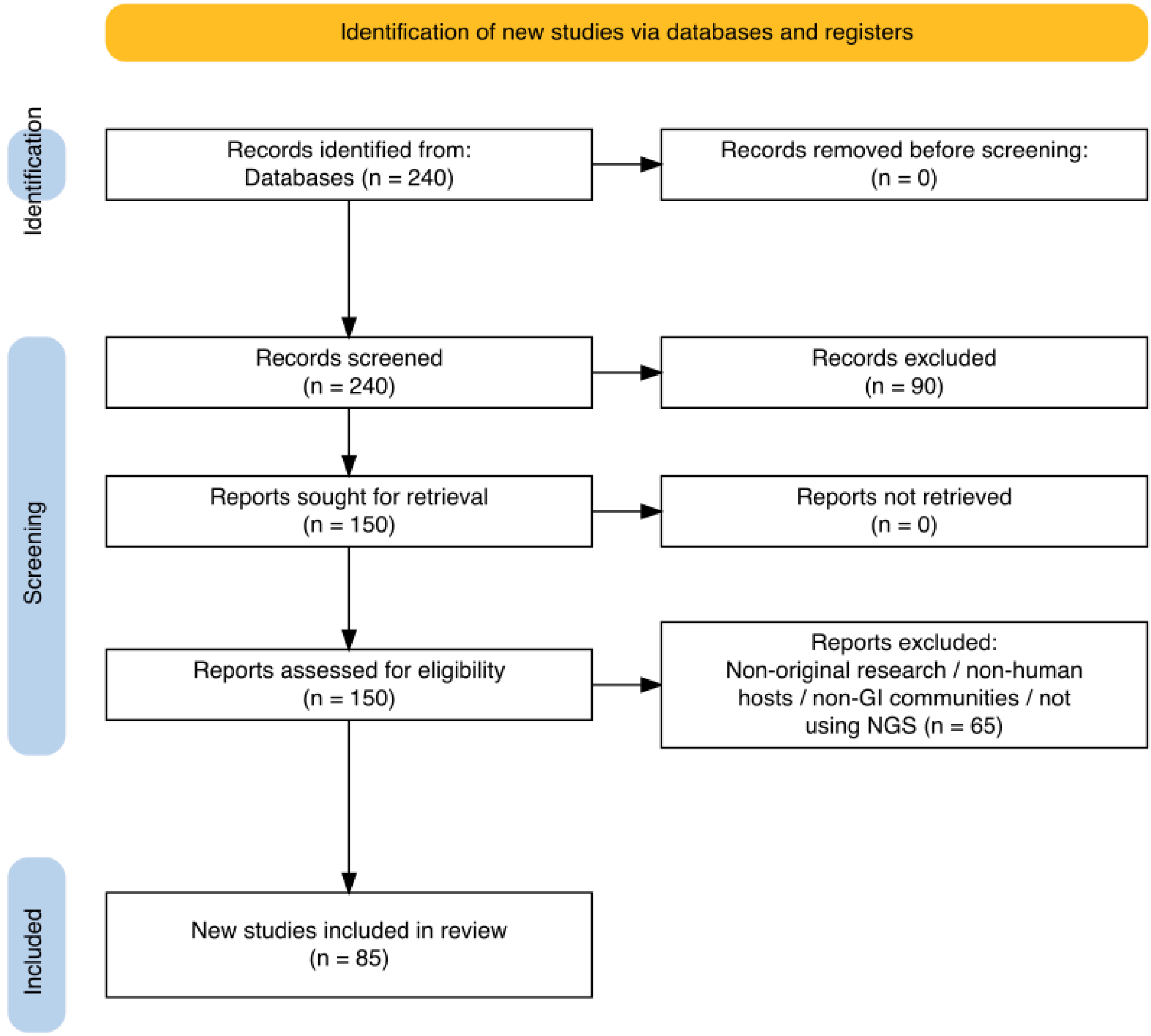


**Supplementary Figure 1.** Schematic of the workflow for selection of publications for Case Study 1. Records were removed during the initial screen (n = 90) for not being available in the PubMed Central Open Access Subset. Diagram was generated with PRISMA2020 (Haddaway, N. R. *et al.* 2020).

The PubMed search query is as follows:

```
((((("journal article"[Publication Type]) NOT ("systematic review"
[Publication Type])) NOT ("review"[Publication Type])) AND
(("2024/01/01"[Date - Publication] : "2024/06/30"[Date - Publication]))) AND
("gut-microbio*"[Title/Abstract] OR "intestinal-microbio*"[Title/Abstract]
OR "gastrointestinal-microbio*"[Title/Abstract])) AND
(human[Title/Abstract])) AND (("amplicon"[Title/Abstract] OR
"16S"[Title/Abstract] OR "16S-rRNA"[Title/Abstract] OR
"marker-gene"[Title/Abstract] OR "metagenom*"[Title/Abstract] OR "shotgun"
[Title/Abstract] OR "nucleotide" [Title/Abstract] OR "DNA"[Title/Abstract])
AND ("sequencing"[Title/Abstract] OR "sequence"[Title/Abstract] OR
"sequences"[Title/Abstract]))
```

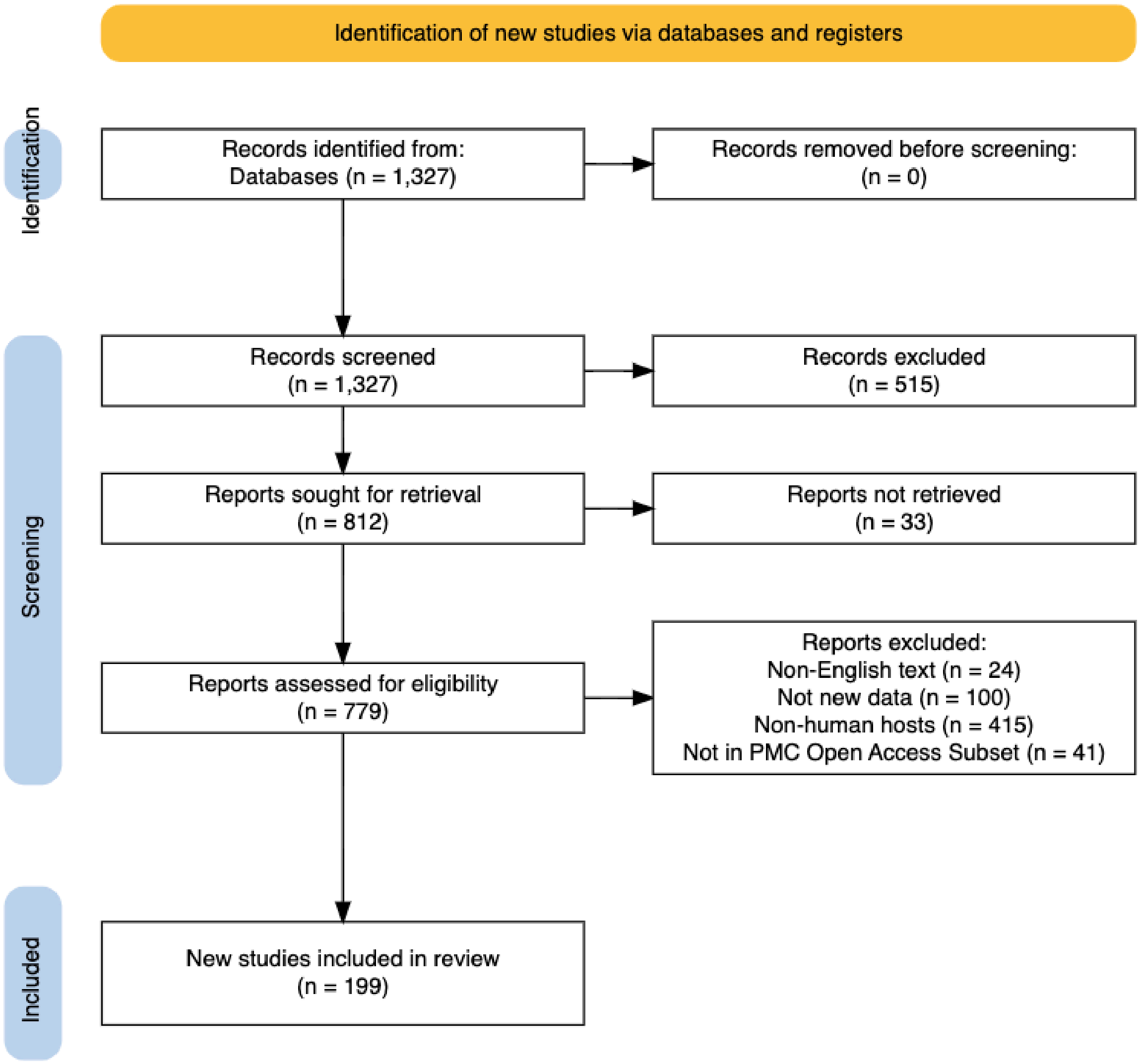


**Supplementary Figure 2.** Schematic of the workflow for selection of publications for Case Study 2. Records were removed during the initial screen (n = 515) for lack of relevance through a manual abstract review. Diagram was generated with PRISMA2020 (Haddaway, N. R. *et al.* 2020).

The PubMed search query is as follows:

```
"intestinal-mycobi* OR gut-mycobi* OR gastrointestinal-mycobi* OR
(intestinal-microbi* OR gastrointestinal-microbi* OR gut-microbi* AND
fung*[tiab]) AND (2003:2023[pdat]) AND journal article NOT (review OR
systematic review)"
```

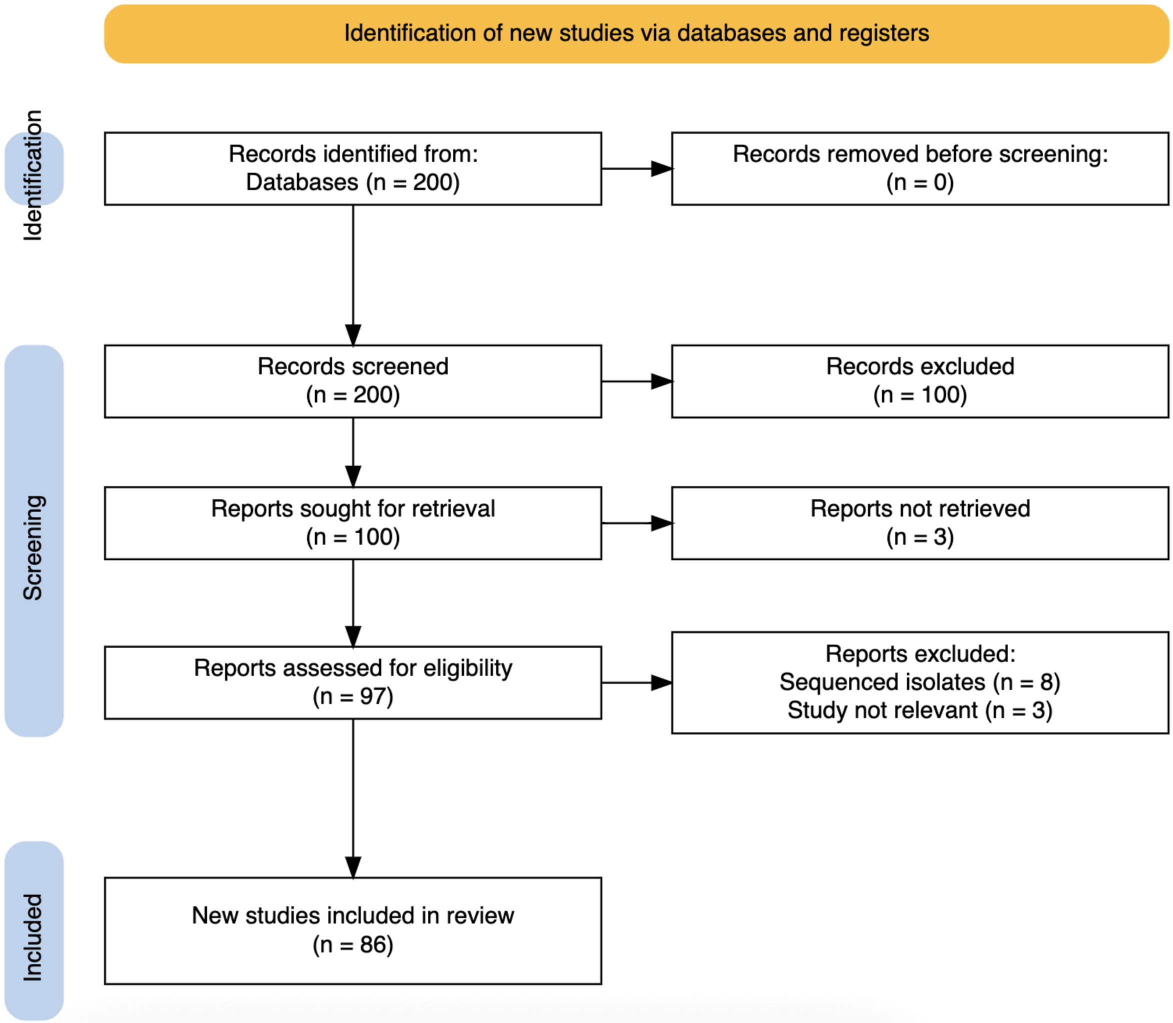


**Supplementary Figure 3.** Schematic of the workflow for selection of publications for Case Study 3. Records were removed during the initial screen (n = 100) for not being available in the PubMed Central Open Access Subset. Diagram was generated with PRISMA2020 (Haddaway, N. R. *et al.* 2020).

The PubMed search query is as follows:

```
soil[Title/Abstract] AND (((("journal article"[Publication Type] NOT
"review"[Publication Type]) NOT "systematic review"[Publication Type]) AND
"soil microbi*"[Title/Abstract] AND "amplicon"[Title/Abstract]) OR
"16S"[Title/Abstract] OR "16S-rRNA"[Title/Abstract] OR
"marker-gene"[Title/Abstract] OR "metagenom*"[Title/Abstract] OR
"shotgun"[Title/Abstract] OR "nucleotide"[Title/Abstract] OR
"sequenc*"[Title/Abstract]) AND 2023/01/01:2023/12/31[Date - Publication]
```

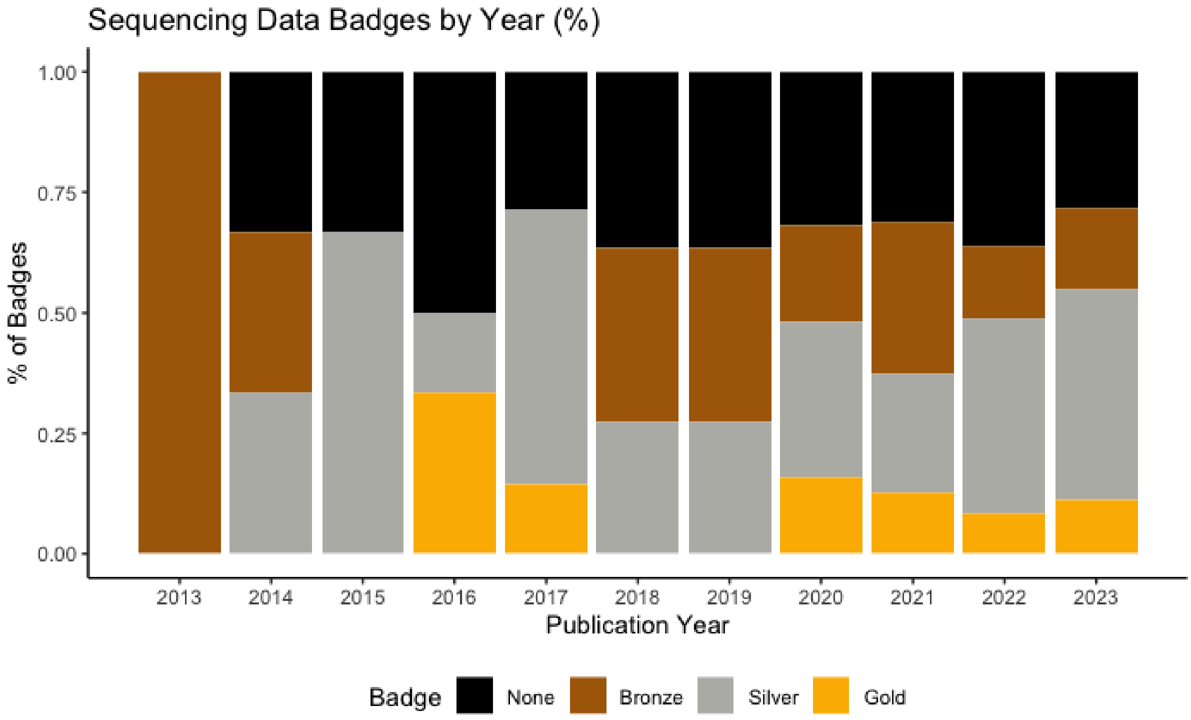


**Supplementary Figure 4.** A temporal look at badge assignments for Case Study 2, human gut-associated mycobiome publications published from 2013 through 2023, by proportion of papers by year.

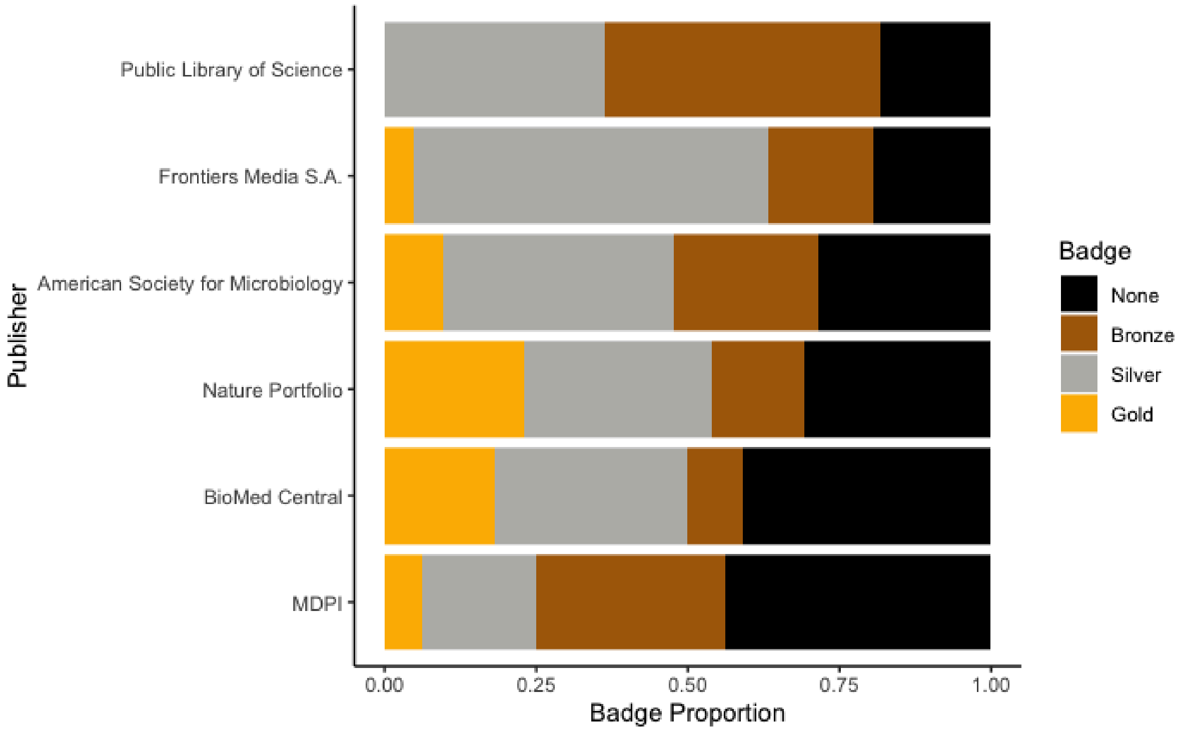


**Supplementary Figure 5.** Badge assignments for Case Study 2, human gut-associated mycobiome publications, categorized by article publishing group. Note that Nature Portfolio and BioMed Central journals, while part of Springer Nature, were sorted into their subsidiaries.
